# EK-DRD: a comprehensive database for drug repositioning inspired by experimental knowledge

**DOI:** 10.1101/515676

**Authors:** Chongze Zhao, Xi Dai, Yecheng Li, Qingqing Guo, Jianhua Zhang, Xiaotong Zhang, Ling Wang

**Author notes:** Equal contributors (Chongze Zhao, Xi Dai, Yecheng Li).

## Abstract

**Summary:** Drug repositioning, or the identification of new indications for approved therapeutic drugs, has gained substantial traction with both academics and pharmaceutical companies because it reduces the cost and duration of the drug development pipeline and it reduces the likelihood of unforeseen adverse events. So far, there has not been a systematic effort to identify such opportunities, in part because of the lack of a comprehensive resource for an enormous amount of unsystematic drug repositioning information to support scientists who could benefit from this endeavor. To address this challenge, we developed a new database, Experimental Knowledge-Based Drug Repositioning Database (EK-DRD) by using text and data mining, as well as manual curation. EK-DRD contains experimentally validated drug repositioning annotation for 1861 FDA-approved and 102 withdrawn small molecule drugs. Annotation was done at four levels, using 70,212 target assay records, 3999 cell assay records, 585 organism assay records, and 8910 clinical trial records. Additionally, approximately 1799 repositioning protein or target sequences coupled with 856 related diseases and 1332 pathways are linked to the drug entries. Our web-based software displays a network for integrative relationships between drugs, their repositioning targets, and related diseases. The database is fully searchable and supports extensive text, sequence, chemical structure, and relational query searches.

**Availability:** EK-DRD is freely available at http://www.idruglab.com/drd/index.php.

## 1 Introduction

The majority of failures of drug development programs are due to the lack of efficacy of therapeutic hypothesis, with unexpected clinical side effects and tolerability being crucial issues (Scannell, et al., 2012). Finding new uses outside the scope of the original medical indication for existing drugs, referred to as drug repurposing or repositioning, is one solution to achieve efficiency. Existing drugs have already been tested in humans, have been demonstrated an acceptable level of safety and tolerability, and are often approved by regulatory agencies for human use (Sachs, et al., 2017). This could potentially increase the success rate of drug development and reduce the cost in terms of time.

Because of the intensifying research and the accumulation of data on drug repositioning, database relevant to drug repositioning has emerged in recent years. PROMISCUOUS is a database that enables users to establish and analyze networks responsible for multi-pharmacology by connecting the measures of structural similarity for drugs and known side-effects to protein–protein interactions (von Eichborn, et al., 2011). Although the above database has aided research into drug repositioning, there is still no specific resource that provides comprehensive data for experimental determination of drug repositioning and further data analysis.

Herein, we developed the Experimental Knowledge-Based Drug Repositioning Database (EK-DRD) to host data on all aspects of experimentally validated repositioning information.

## 2 Methods

### 2.1 Data collection and processing

A total of 1963 small molecule drugs were retrieved from DrugBank version 4.0 (Law, et al., 2014). First, the drugs with available experimentally determined target assay data were searched from the public databases of ChEMBL (Bento, et al., 2014), BindingDB (Gilson, et al., 2016), PubChem BioAssay (Wang, et al., 2014), and PDSP Ki. The target assay data were refined with the criteria as the follows: (1) only target-based assay data with detailed assay values (e.g., *K_i_* or IC_50_) were kept; (2) the ADMET assay data were excluded; (3) the repositioning target assay data were filtered and obtained by mapping the FDA approval target. Second, the cell-based assay data for repositioning were obtained by searching ChEMBL. Third, the organism-based assay data were retrieved from PubMed using the combinations of the keywords “drug name and synonyms”, “*in vivo*”, “organism”, and “animal”. The searched literatures were evaluated to find drugs with repositioning datasets. Finally, the repositioning data for clinical trial indications were obtained from the American Association of Clinical Trials Database. All public databases we accessed were up to September 2016. All of these repositioning data for 1963 drugs from different sources were checked by manual curation. Related information for drug repositioning, such as repositioning targets (gene, function, sequences, structures, etc.), signal transduction pathways, and diseases, were retrieved from UniProt (Bateman, et al., 2015), PDB (Rose, et al., 2017), KEGG (Kanehisa, et al., 2017), and TTD (Yang, et al., 2016) databases.

### 2.2 Database and web interface implementation

All of the metadata are stored and managed in a MySQL database. The web interfaces were implemented in HTML, JavaScript and PHP. The ChemDoodle web component (web.chemdoodle.com/) is used as a 2D molecule editor. Three retrieval methods (text, chemical structure, and protein sequence searches, Part I of the Supplementary Material) are provided for users to access the EK-DRD. Technologies for creating EK-DRD are summarized in Supplementary Table S1.

## 3 Results and discussion

### 3.1 Database content and statistics

EK-DRD contains 1963 small molecule drugs with four different types of repositioning bioassay data: target, cell, organism, and clinical trials (Supplementary Fig. S1). For the target level, there are 70,212 assay data points for 1799 different repositioning targets from 123 different species. Approximately 89.22% of the repositioning targets come from the top nine species (Supplementary Fig. S2A). For the cell level, 3999 repositioning assay data points were stored in EK-DRD. For the clinical trial level, 8910 clinical trials were annotated for drug repositioning. The proportions for repositioning for different stages of clinical research are shown in Supplementary Fig. S2B.

EK-DRD contains many associated data (Table S2), such as information related to 1799 repositioning targets (gene, function, sequences, structures, etc.), 1332 signal transduction pathways, and 856 related diseases involved in these repositioning targets, comprising 3762 drug–repositioning target–disease networks that are drug-centric and repositioning-target-centric.

### 3.2 Web interfaces and their usage

Detailed description of web interfaces and their usage can be found in Part II of the Supplementary Material and Fig. S3, S4 and S5.

## 4 Conclusions

With the growing number of drug-repositioning studies, there is need for an integrated database that facilitates the exploration of data from these studies. A comparison of EK-DRD and PROMISCUOUS is provided in Supplementary Table S3. To the best of our knowledge, EK-DRD is the first publicly available comprehensive resource for hosting and analyzing experimental-knowledge-based drug-repositioning datasets. The expanded coverage of experimentally validated drug repositioning data, together with the knowledge of the mechanisms, chemical structures and properties of drugs as well as drug-repositioning target-disease networks, can facilitate repositioning-based drug discovery and related development, optimization, or both of *in silico* tools.

