## Supplemental Figure1-5 Table 1-3 for "EK-DRD: a comprehensive database for drug repositioning inspired by experimental knowledge"

†Equal contributors. *To whom correspondence should be addressed.

**Part I: Search and network-display tools**

EK-DRD provides three retrieval methods for quickly searching and displaying the drug repositioning data, namely, text mining, chemical structure search, and protein sequence search. For chemical structure search, five algorithms, namely, substructure search, Markush search, two-dimensional (2D) and three-dimensional (3D) similarity calculations, and hybrid structure-similarity calculations, are used in EK-DRD. 2D similarity calculations are based on the FP2 fingerprint and performed using OpenBabel (O'Boyle, et al., 2011). 3D similarity adopts the weighted Gaussian algorithm (WEGA) for molecular-shape-similarity calculations (Yan, et al., 2013), which provides shape-, feature- and coefficient-based shape-feature combo-scoring functions for user selection. Our group also encoded in EK-DRD a new hybrid-similarity metric for calculating compound similarity that combines 2D fingerprint and 3D shape, called HybridSim, which was developed and validated to outperform the popular 2D FP2-, MACCS-, and 3D WEGA-based similarity methods (Shang, et al., 2017). All similarity methods use Tanimoto coefficient as a similarity function to quantify the similarity between two molecules. BLAST algorithm is used for protein sequence similarity search (Altschul, et al., 1990). We also developed an online network-display tool to virtually display the relationship among drugs, repositioning putative protein targets, and related diseases, in the form of an interactive network.

### Part II: Web interfaces and their usage

EK-DRD provides fast, versatile, and user-friendly web interfaces that enable users to search, browse, display, and download all of the experimentally obtained drug-repositioning data in the database. Moreover, the Contribute data module in EK-DRD can be used to add new drug repositioning data from public users and researchers in the field.

***Search.*** EK-DRD provides three modes of query of the database, i.e., keywords (drug name, drug CAS number, target name, and Uniprot ID), chemical similarity to the EK-DRD drug entries, and sequence similarity to EK-DRD target entries. Here, we present a 2D similarity chemical search as an example to show how to utilize EK-DRD through the search function (Supplementary Fig. S3A). We sought drug repositioning information on dasatinib (a cancer drug). Using ChemDoodle (http://www.chemdoodle.com) sketcher, a user can build a molecular structure of dasatinib and click the button of "Search By Draw" to perform the 2D similarity search with other drugs in the EK-DRD database. The 2D similarity search results are ranked by similarity score and shown in Supplementary Fig. S3B. The first record is dasatinib, which has the highest similarity score of 1; the users can click "Show Detail" button to enter the repositioning information page of dasatinib (Supplementary Fig. S3C), which displays basic information, FDA approval induction and target, as well as repositioning data at target, cell, organism, and clinical trial levels. The user may check and browse the detailed data for target-level assays (Supplementary Fig. S3D). In addition, the connection concept network of dasatinib-repositioning target-related disease (Supplementary Fig. S3E) can be found in the repositioning information page.

***Browse.*** Repositioning information for drugs and targets can be browsed in two ways: (1) Users can directly find the detailed repositioning data of the desired drugs according to drug name in alphabetical order (e.g., abacavir, Supplementary Fig. S4A); and (2) According to alphabetical order of repositioning target name, the repositioning target page displays basic information on the desired target (target name, gene name, PDB ID, KEGG ID, pathway ID, and repositioning drugs) and associated descriptions of functions, related diseases, and pathways as well as the hyperlink for repositioning-target-centric network (e.g., A7 nicotinic acetylcholine receptor, Supplementary Fig. S4B).

***Network.*** This page displays drug–target–disease networks that are drug-centric and repositioning-target-centric. For the drug-centric-based network (e.g., dasatinib, Supplementary Fig. 5A), which is based on experimentally determined repositioning target–drug interactions, the repositioning target may be involved in specific physiological functions for the treatment of certain diseases. Therefore, linking the drug and repositioning targets to their treated diseases is highly useful; this suggests that the repositioning-target-related diseases may be potentially treated by the desired drug. On the basis of this view, we built a such drug-centric-based drug–target–disease network. The repositioning-target-centric-based network (e.g., A7 nicotinic acetylcholine receptor, Supplementary Fig. 5B) links the target and related diseases to all repositioning drugs, thus helping users to check the potential combination therapy. Users can browse the drug–target–disease network in terms of drug or target. In drug–target–disease network, the users can double click on the identifier of the desired target or drug to browse their detailed information. In addition, the Search module (input drug, target name and Uniprot ID) in the Network page enables users to find the drug–target–disease network of the desired drug or repositioning target.

***Contribute Data.*** If public users and researchers know of or have new experimentally determined drug repositioning data that they would like us to add, they can download the template table (CSV format) for the target, cell, organism, and clinical trial at the "Contribute Data" page. They can fill the table and send it to us using the Submit module at the "Contribute Data" page.

***Download and FAQ.*** All data in the EK-DRD database can be freely downloaded from the "Download" page, and a detailed introduction and tutorial on the EK-DRD database are available on the "FAQ" page.

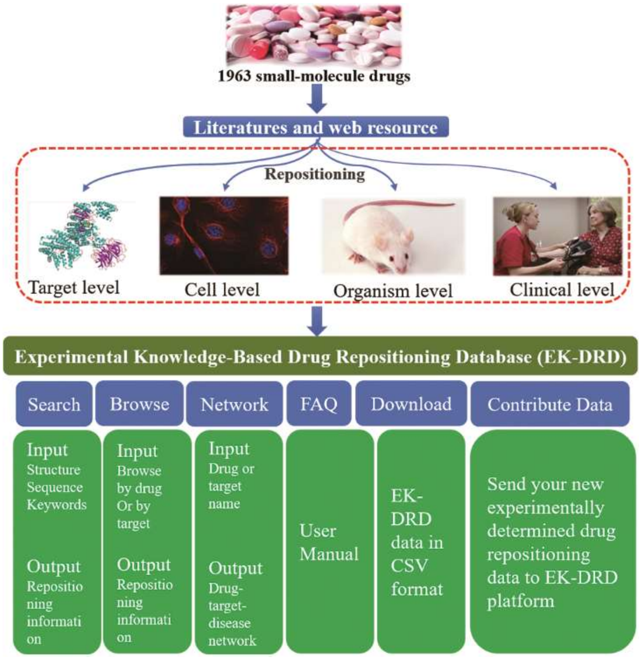


**Supplementary Fig. S1.** Overall design, construction, and contents of EK-DRD.

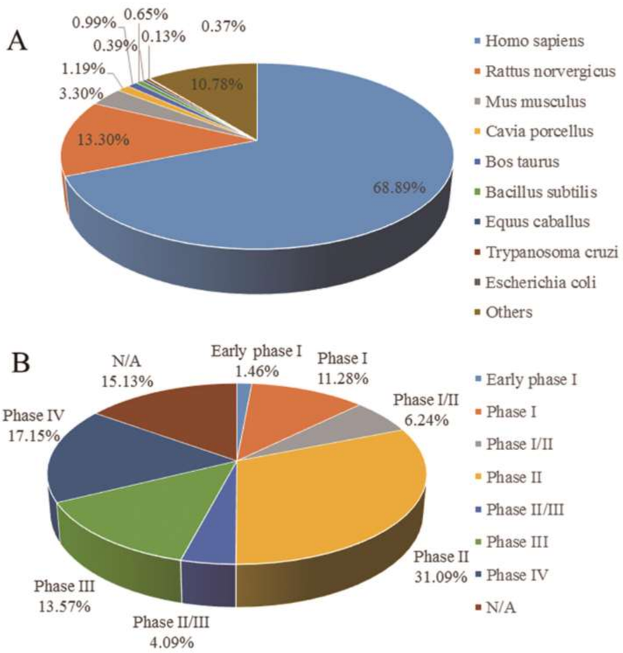


**Supplementary Fig. S2.** Distribution of repositioning targets for 1963 drugs among the different species (**A**) and repositioning clinical trial records for these drugs (**B**) in EK-DRD. N/A (not applicable) is used to describe trials without FDA-defined phases, such as trials of devices or behavioral interventions.

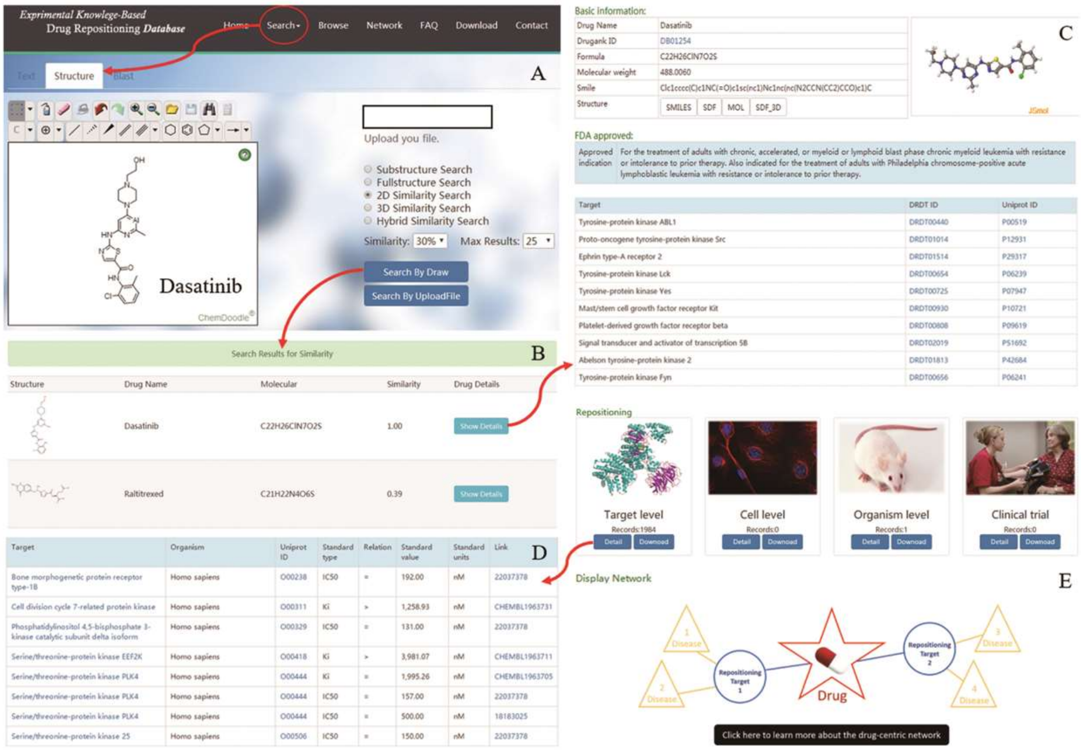

**Supplementary Fig. S3.** A schematic workflow of the chemical structure search interface in EK-DRD. (**A**) 2D chemical similarity for dasatinib drawn by using the online ChemDoodle sketcher. (**B**) Snapshot of search results for dasatinib obtained by using the 2D similarity search mode. (**C**) Snapshot of basic, FDA-approved, and repositioning information on dasatinib. (**D**) Detailed target-based repositioning information for dasatinib presented as a table. (**E**) The connection concept network of dasatinib-repositioning target-related disease.

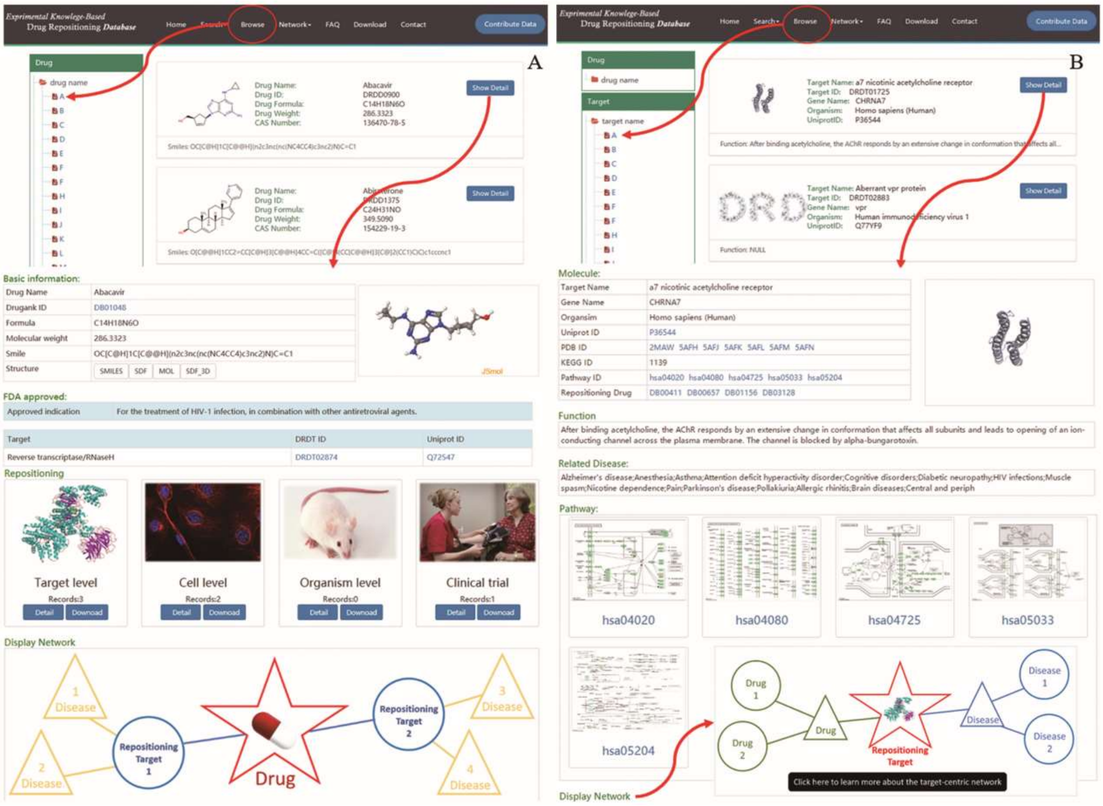

**Supplementary Fig. S4.** Repositioning information for drugs and targets can be browsed in alphabetical order according to drug or target name. **(A)** Snapshot of the detailed repositioning data of abacavir. **(B)** Snapshot of the repositioning target page for A7 nicotinic acetylcholine receptor.

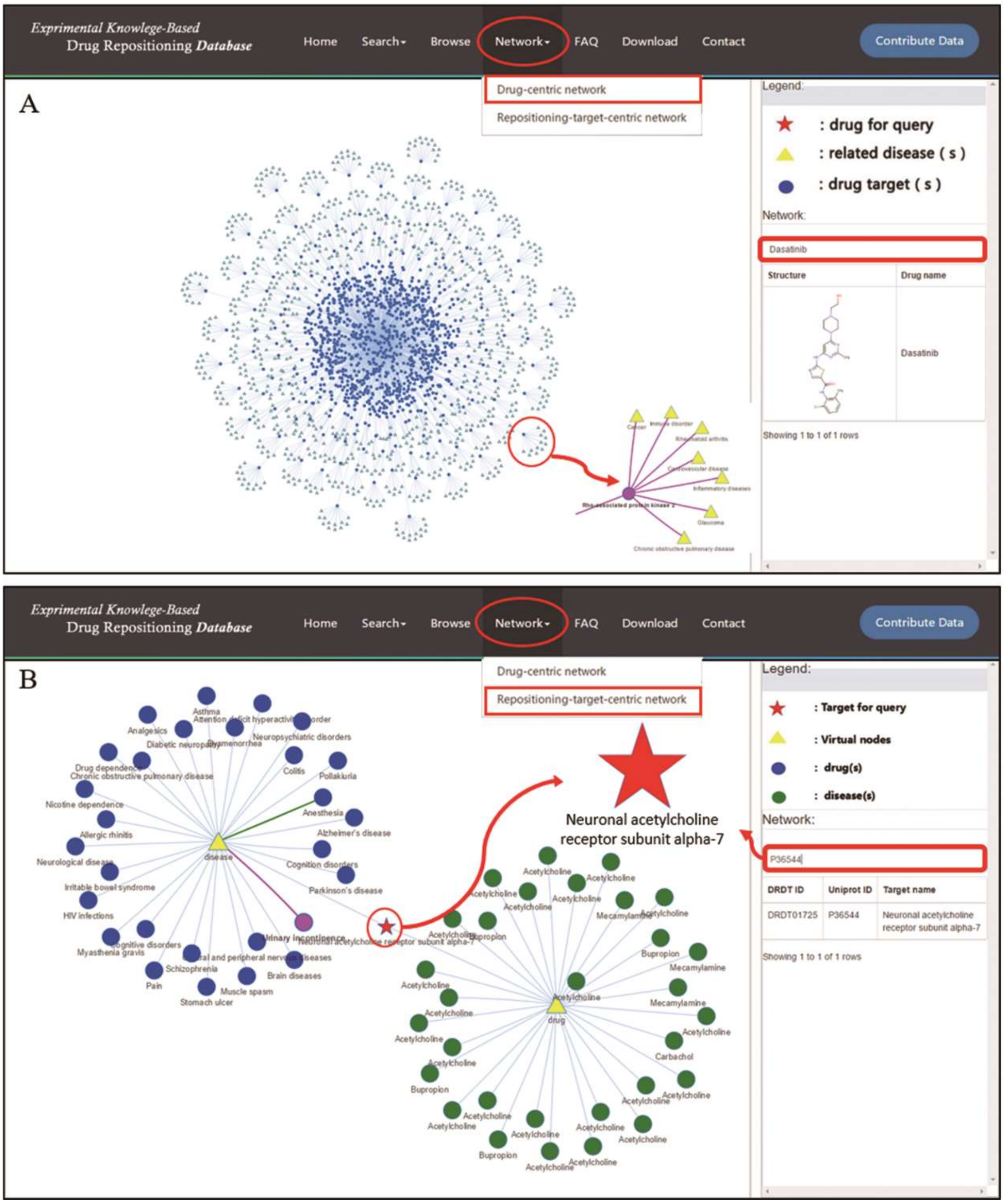


**Supplementary Fig. S5.** The drug–repositioning target–disease networks web page. (**A**) Snapshot of the drug-centric network for dasatinib. (**B**) Snapshot of the repositioning-target-centric network for A7 nicotinic acetylcholine receptor.

**Supplementary Table S1**. The toolkits used to create the EK-DRD database.

| Tools/Database | Purpose | Source |
|---|---|---|
| ChemDoodle Web | structure draw online | web.chemdoodle.com/ |
| Django | a high-level Python Web framework | https://www.djangoproject.com/ |
| NGINX | an HTTP and reverse proxy server, a mail proxy server, and a generic TCP/UDP proxy serve | http://nginx.org/en/ |
| Open Babel | uniformly converted all drug structures in multiple formats | http://openbabel.org/wiki/Main_Page/ |
| WEGA | 3D shape-based similarity calculation | *In-house* |
| HybridSim | Molecular-similarity searches based on 2D fingerprint and 3D shape | http://www.idruglab.com/HybridSim-VS/ |
| MOE software | Generate 3D structure for each drug | http://www.chemcomp.com/ |
| Discovery Studio software | Conformational ensembles generated for each drug in the database | http://accelrys.com/products/collaborative-science/biovia-discovery-studio/ |
| PubMed | PubMed is a database that provides biomedical articles | https://www.ncbi.nlm.nih.gov/m/pubmed/ |
| ClinicalTrials.gov | American Association of Clinical Trials Database | https://clinicaltrials.gov/ct2/ |
| Drugbank | US FDA approval drugs source (only for small molecules) | https://www.drugbank.ca/ |
| ChEMBL | manually curated chemical database of bioactive molecules with drug-like properties | https://www.ebi.ac.uk/chembl/ |
| BindingDB | curated measured binding affinities | https://www.bindingdb.org/ |
| PubChem BioAssay | a biochemical experimental database | https://www.ncbi.nlm.nih.gov/pcassay/ |
| PDSP Ki | a database of ligand and target affinity | https://pdsp.unc.edu/databases/kidb.php/ |
| UniProt | UniProt is a freely accessible database of protein sequence and functional information | http://www.uniprot.org/ |
| PDB | a database of biological macromolecular structures | https://www.rcsb.org/ |
| TTD | a database to provide information about the known and explored therapeutic targets | https://db.idrblab.org/ttd/ |
| MySQL | manage and store all of the metadata | http://www.mysql.com |
| HTML | a language for describing web documents | https://tutorialehtml.com/ |
| CSS | a style sheet language used for describing the presentation of a document written in a markup | https://www.w3.org/Style/CSS/ |

| | | |
|---|---|---|
| | language | |
| Apache HTTP server | maintain an open-source HTTP server for modern operating systems | http://http.apache.org/ |
| JavaScript | a high-level, interpreted programming language | https://www.javascript.com |
| PHP | a popular general-purpose scripting language that is especially suited to web development | http://www.php.net/ |

**Supplementary Table S2.** Summary of the data fields or types and primary drug repositioning records in EK-DRD.

| Drug information | Target/Cell/Organism/Clinical trial information | Num. of repositioning records and networks |
|---|---|---|
| Common name | Target name | Target level (70,212) |
| Chemical structure | Gene Name | Cell level (3999) |
| CAS number | Organism | Organism (585) |
| Chemical formula | Uniprot ID link | Clinic trial (8910) |
| DrugBank/ChEMBI ID Links | KEGG/Pathway ID links | Drug-centric based drug-target-disease network (1963) |
| Molecular weight | Target Function | Target-centric based drug-target-disease network (1799) |
| Chemical formula | Related disease | |
| SMILES string | Target pathway | |
| MOL file | Target PDB ID | |
| SDF (2D/3D) files | Target sequence | |
| FDA approved indication | Cell name | |
| Mechanism of action | Cell-based assay description | |
| | Standard assay type/value/units | |
| | Organism-based assay description | |
| | Clinical trial basic description (Title, Interventions, Phase, Disease) | |
| | ClinicalTrials.gov Identifier link | |

**Supplementary Table S3.** The differences of EK-DRD and PROMISCUOUS.

| Data/Feature | EK-DRD | ROMISCUOUS |
|---|---|---|
| Drugs | 1963 (FDA approval and withdrawn) | 25000 (FDA approval, withdrawn or experimental drugs) |
| Data source | Drugbank;<br>ChEMBL;<br>BindingDB;<br>PubChem BioAssay;<br>PDSP Ki;<br>PubMed | Drugbank;<br>SuperTarget;<br>SuperCyp;<br>PubMed |
| Data Type | Experimentally determined | Experimentally determined and/or inferred relationships through structural similarity |
| Target level assay data | Yes, 70,212 assay records,<br>with detailed assay description and values | Yes, 21,500 drug-protein interactions relationships, without detailed assay description and values |
| Cell level assay data | Yes, 3999 assay records,<br>with detailed assay description and values | No |
| Organism assay data | Yes, 585 assay records,<br>with detailed assay description | No |
| Clinical trial data | Yes, 8910 clinical trial records,<br>with detailed description | No |
| Text search | Yes | Yes |
| Chemical structure search | Substructure: Yes<br>2D similarity: Yes, configurable settings<br>3D similarity: Yes, configurable settings<br>Hybrid-similarity: Yes, configurable settings | Substructure: No<br>2D similarity: Yes, unconfigurable settings<br>3D similarity: No<br>Hybrid-similarity: No |
| Sequence search | Yes | No |
| Network analysis | Yes, drug-repositioning target-disease network (experimental data-driven) | Yes, drug-target side effects network (computational similarity data-driven) |
